# The nose knows: Nasal temperature tracks facial attractiveness, not social categorization

**DOI:** 10.64898/2026.01.26.701747

**Authors:** Erik Van der Burg, Ivo V. Stuldreher, Tim Ziermans

## Abstract

Facial attractiveness plays a central role in first impressions, social interactions, and romantic relationships, yet remains difficult to quantify objectively due to its subjective and socially shaped nature. In the present study, we examined whether facial attractiveness and its modulation by social information can be captured using functional infrared thermal imaging (fITI). Participants rated the attractiveness of faces that were randomly preceded by an autism label. Although such labels influenced explicit attractiveness judgments, particularly among male participants, they did not modulate facial thermal responses. Instead, nose temperature systematically increased or decreased when participants rated faces as attractive or unattractive, respectively. Notably, temperature differences emerged several seconds after image onset, and for female faces, mean attractiveness ratings positively correlated with changes in nose temperature. Together, these findings reveal a dissociation between socially shaped explicit evaluations and autonomic physiological responses, highlighting the potential of fITI as a fully non-invasive tool for capturing implicit affective engagement with facial attractiveness.

---

Facial attractiveness plays a central role in shaping first impressions, guiding social interactions, and influencing romantic choices. What counts as attractive can be operationalized through well-established facial cues such as averageness, skin colour, masculinity, symmetry and more (Langlois & Roggman, 1990; Rhodes, 2006; Rhodes et al., 1998, 2001; Thornhill & Gangestad, 1999). Meta-analytic evidence reveals striking agreement in attractiveness judgments both within and across cultures (Langlois et al., 2000), challenging the idea that beauty standards are merely cultural artefacts. At the same time, considerable research shows that personal experiences, individual differences, and social biases also shape how people evaluate facial attractiveness (see Little et al., 2011, for a review). In the present study, we examine whether autism-related stigma affects perceived facial attractiveness and whether such biases can be captured through objective physiological responses using functional infrared thermal imaging (fITI).

A recent study by Brosnan and Gavin (2021) investigated whether autism stigma influences judgments of attractiveness when women evaluated a man’s dating profile. Furthermore, they examined whether participants’ level of stigmatization of, or familiarity with, autism moderated these evaluations. Interestingly, the authors found that the presence of an explicit autism label increased perceived attractiveness relative to an unlabeled dating profile (see also Gavin et al., 2019). Although stigmatization and familiarity did not directly alter attractiveness ratings, lower levels of stigma were associated with a greater desire to date the profiled individual. In contrast, Aydin and colleagues (2024) reported that an attractive male face was judged as less attractive when labeled as having schizophrenia, suggesting that diagnostic labels can bias social evaluations in condition-specific ways. Taken together, these findings demonstrate that diagnostic labels powerfully shape perceived attractiveness but also underscore the limits of relying exclusively on self-reported judgments. This highlights the importance of incorporating objective physiological measures that can detect evaluative biases independent of conscious self-report.

While diagnostic labels can clearly bias attractiveness evaluations, it remains unclear whether such biases affect only what people consciously report or also their automatic physiological responses. fITI offers a sensitive and non-invasive method for capturing these responses by measuring facial temperature fluctuations associated with sympathetic and parasympathetic activity. Thermal patterns in regions such as the nose, forehead, and areas around the eyes (periorbital regions) are known to reflect affective and evaluative processing, including sexual arousal, fear, joy, and guilt (see Ioannou et al., 2014, for a review). For example, Hahn and colleagues (2012) demonstrated that nasal temperature increased when participants were touched on highly intimate body locations (face and chest) compared to less intimate locations (arm and palm), with effects strongest during opposite-sex contact, highlighting the sensitivity of facial thermal responses to socially evaluative and affective contexts (see also Merla & Romani, 2007, for a similar finding). Despite this growing body of work, it remains unknown whether facial thermal responses are sensitive to perceived facial attractiveness perse. Because these physiological changes occur largely outside conscious control, fITI provides an implicit index of evaluative processing that complements traditional rating-based measures.Combining fITI with subjective judgments therefore enables a more comprehensive assessment of how diagnostic labels influence the perception of facial attractiveness.

The present study aimed to test whether an autism label biases facial attractiveness evaluations and, critically, whether such bias is expressed only in explicit judgments or also in automatic physiological responses measured with fITI. Participants viewed a series of either 48 male or female faces, each preceded by a randomly assigned label indicating either “person with autism” or “person without autism,” and rated the attractiveness of each face using a rating scale (see also, Kok et al., 2017; Taubert et al., 2016; Van der Burg et al., 2019). Figure 1a illustrates an example trial.Concurrently, facial thermal responses were recorded from the tip of the nose, a region shown to be particularly sensitive to affective and social–evaluative processing (see Figure 1b for an example thermal image). Based on previous findings (Brosnan & Gavin, 2021), we hypothesized that faces labeled as autistic would be judged as more attractive than presented without such a diagnostic label. In addition, drawing on prior work showing that nasal temperature changes track affective and social arousal (Hahn et al., 2012; Ioannou et al., 2014; Merla & Romani, 2007), we tested whether facial attractiveness is associated with systematic changes in nasal temperature. Specifically, we predicted that higher perceived attractiveness would be accompanied by increased nasal temperature, whereas lower attractiveness would be associated with decreased temperature. By combining subjective ratings with physiological measures, this design allows us to determine whether autism-related stigma influences only explicit evaluations or also the underlying perceptual and affective processes accompanying judgments of facial attractiveness. To further explore individual differences, participants completed two brief questionnaires assessing stigma toward and familiarity with autism, allowing us to test whether these factors moderated both attractiveness ratings and physiological responses. Finally, we examined whether individual differences in empathy predicted attractiveness judgments and nasal temperature changes, given evidence that empathy modulates autonomic and neurophysiological responses during social perception (Almeida et al., 2024; D. Neumann & Westbury, 2011).

**Figure 1.**
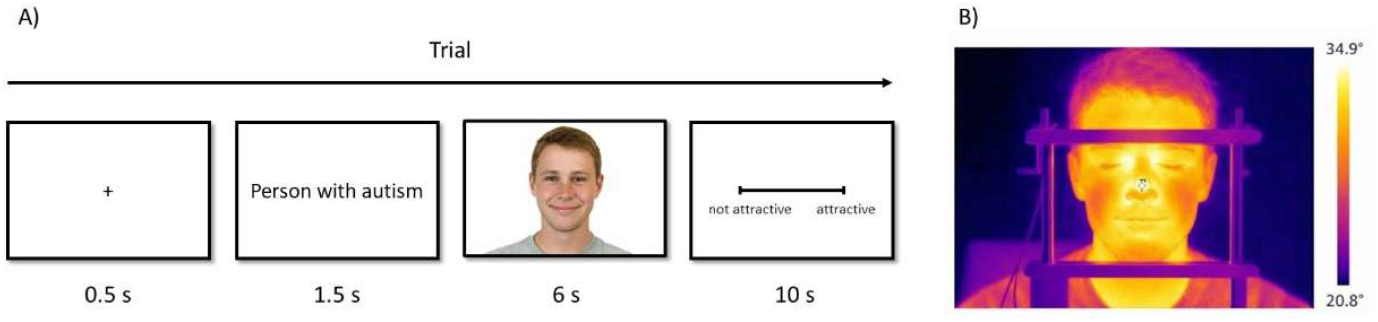
Experimental design and thermal recording. A) Example trial. Participants first viewed a fixation cross, followed by a blank screen displaying a label (“person with autism” or “person without autism”). Next, a facial image was presented for 6 s, after which participants rated its attractiveness on a continuous scale from 0 to 100 using a mouse within a 10 s response window. B) Example thermal infrared image of a participant positioned in a chinrest during the task. The faces shown in this figure are AI-generated images used for illustrative purposes; no real individuals are depicted.

## Method

### Participants

Forty-six volunteers participated in the present study (32 females, 14 males; mean age was 21.8 years, ranging from 18 to 28 years). None of the participants reported having a formal diagnosis of autism, or any other psychiatric condition. The participants were naive to the purpose of the experiment and signed an informed consent prior to the experiment. The experiment was approved by the Ethics Review Board of the Psychology department at the University of Amsterdam (FMG-2561), and in accordance with the Helsinki Declaration.

### Stimuli and apparatus

To measure empathy, participants completed the Dutch validated, shortened version (Carré et al., 2013) of the Basic Empathy Scale (BES; Jolliffe & Farrington, 2006), consisting of 20 items to assess cognitive (9 items) and affective (11 items) empathy (see, Cabedo-Peris et al., 2021 for a review). Participants were asked to rate the degree to which they agreed or disagreed with each statement using a 5-point Likert scale (ranging from 1: totally disagree to 5: totally agree). Participants also completed a questionnaire on their level of stigmatization of, and familiarity with, autism. An adapted version of the Social Distance Scale (SDS; Bogardus, 1933) was used to assess stigma towards individuals with autism (Gillespie-Lynch et al., 2015). The questionnaire consists of six items assessing how willing participants are to interact with a person with autism at increasing levels of social proximity. Participants rated the extent to which they agreed or disagreed with each statement on a 4-point Likert scale (1 = least stigma, 4 = most stigma). Total scores ranged from 6 to 24, with higher scores indicating greater stigma toward autism. Another questionnaire was used to quantify participants’ familiarity with, or level-of-contact with, autistic individuals (Gardiner & Iarocci, 2014). The questionnaire consists of 12 statements (in random order) describing varying levels of exposure to a person with autism, each assigned a relative rank.Participants were instructed to select all items that applied to their own experience, and their score was determined by the rank of the item representing the most intimate level of contact (from “I have never observed a person that I was aware had autism” to “I have autism”). For example, if a participant endorsed the highest-ranked item, “I have autism,” they would receive a score of 12, regardless of the other items selected.

The experiment was programmed in Python using the Psychopy module (Peirce *et al*., 2019). Participants used a chinrest to minimize motion and sat at approximately 50 cm from a 12-inch Dell Latitude E7270 laptop (60 Hz refresh rate) to perform the task using a standard optical mouse. We randomly selected 50 different male and 50 different female colour photos (72% white faces, and 28% black faces) with a mean age of 26.6 years (SD=4.66) from the Chicago Face Database (Ma, Correll, & Wittenbrink, 2015). All photos (2444 pixels width x 1718 pixels height) were taken from the front view under symmetric and uniform lighting conditions. The distance between the camera and the face was kept constant, and the emotional expression of each face was happy. The face images were presented at the centre of the screen (see Figure 1 for an illustration). The background was white and kept constant during the experiment. The rating scale consisted of a centrally presented black line, and both ends were capped with short black lines to signify the upper and lower limit of the scale. The words ‘not attractive’ and ‘attractive’ were shown below the left and right end of the scale, respectively (see also Taubert, Van der Burg, & Alais, 2016; Van der Burg, Rhodes, & Alais, 2019). Participants could click along this line to confirm their rating (on a scale ranging from 0-100).Thermal images were recorded using a FLIR T660 infrared camera with a 7.5 Hz sample rate (resolution: 640×480 pixels). The distance between the infrared camera and the participant was fixed at 110 cm.

### Design and procedure

Participants acclimatised for a duration of 15 minutes. During this period, the participants completed the BES. Subsequently, the participants started with the experiment and indicated whether they wanted to rate the attractiveness of male or female faces during the experiment (13 out of 14 males rated female faces, and 29 out of 32 females rated male faces). Each trial started with the presentation of a black fixation cross for 500 ms, followed by a label “Person with autism” or “Person without autism” for 1.5 seconds. Half of the face images were labelled as autistic, whereas the remaining faces were labelled as non-autistic. The order of the label and type were randomly determined on each trial. Subsequently, participants saw a face image for 6 seconds, followed by the rating scale, and instructed to rate the attractiveness of the face by using the mouse pointer within 10 seconds interval (the screen became blank when the participants made their response). The next trial was initiated after these 10 seconds. The experiment consisted of a single experimental block of 48 trials. Participants received instructions prior to the experiment and performed two practice trials to become familiar with the task and procedure. After the experiment, participants completed the two questionnaires to measure i) familiarity with autism, and ii) the SDS to measure the willingness to join people with autism.

### Analyses

Statistical analyses were conducted using Just Another Statistical Program (JASP; Love *et al*., 2019) or Python (version 3.11.14). For all analyses, α was set to.05. For each participant, we calculated the median attractiveness rating over all the faces and labelled each face as attractive if the rating was greater than the median rating, or as unattractive if the rating was equal to, or lower than the median. The statistical tests conducted are mentioned at the appropriate locations in the Results section.

For each trial, we measured the nose temperature (a single pixel) from the face image onset for 16 seconds (i.e., 120 samples given a 7.5 Hz sample frequency). Subsequently, the nose temperature was baseline corrected by subtracting the temperature at the stimulus onset so that we measured the nose temperature change relative to the temperature at stimulus onset (instead of measuring the absolute temperature). Then, for each trial we determined whether participants moved or not. Trials were excluded from further analyses if the temperature changed more than 2.5 ºC in either direction from the trial onset at a given sample.

## Results

Data from six participants were removed from further analyses. Three participants were excluded as their mean attractiveness rating was too close to zero (mean score was 3.5 on a scale from 0-100; group mean was 35.1). Three other participants were excluded as they continuously moved their head during the experiment (100% of the trials; group average was 3.98%). The two practice trials and trials on which participants moved their head (3.98%) were excluded from further analyses. Figure 2 illustrates the results of the remaining 40 participants.

**Figure 2.**
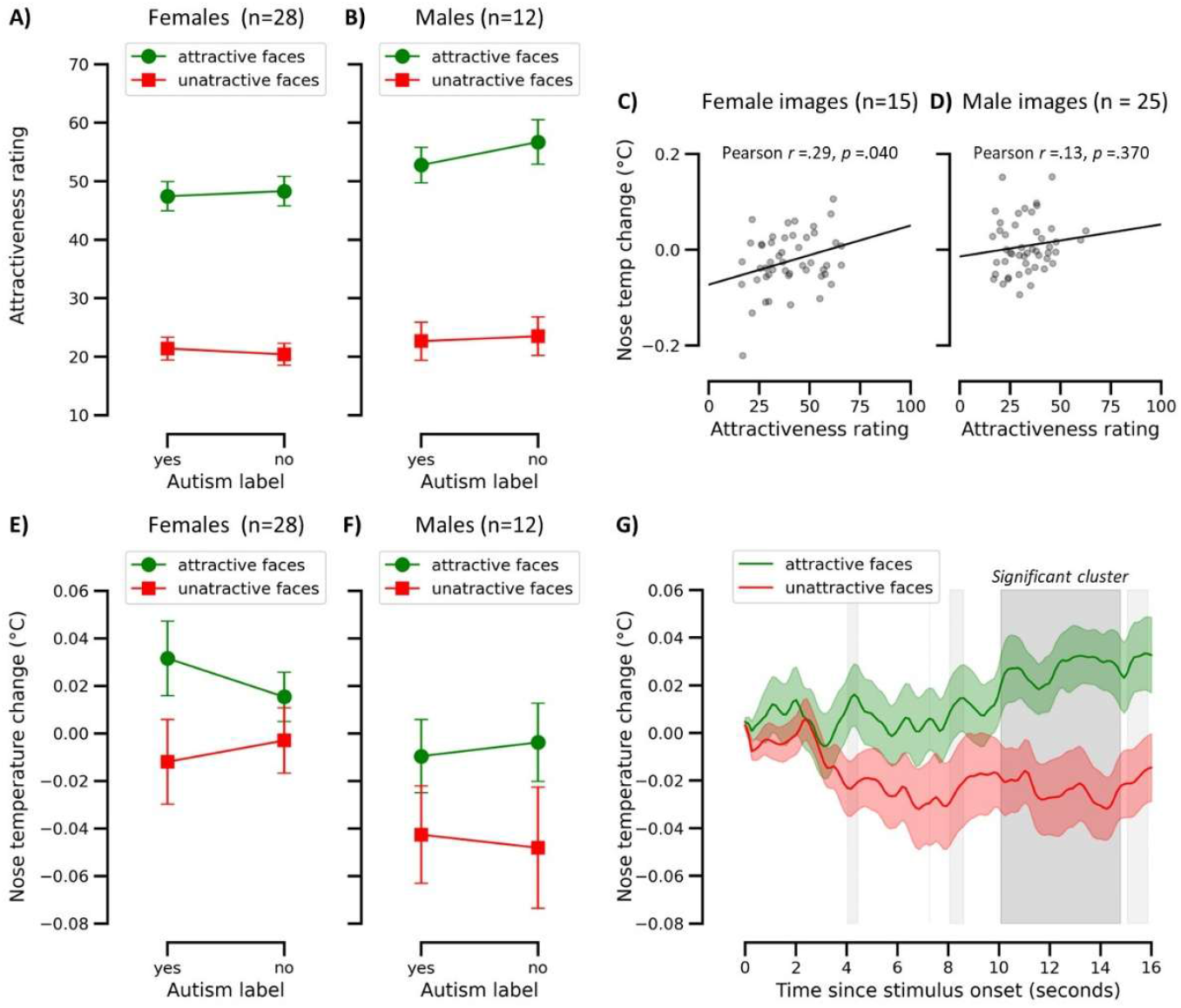
A and B) Mean attractiveness rating as a function of autism label and facial attractiveness for female (Panel A) and male participants (Panel B). C and D) Correlation between the mean nose temperature change and attractiveness rating for female (Panel C) and male images (Panel D). Note that the 48 female images were rated by 12 males and 3 females, whereas 48 male images were rated by 25 females. E and F) Mean nose temperature change as a function of autism label and facial attractiveness for female (Panel E) and male participants (Panel F). G) Time course of the mean nose temperature change as a function of facial attractiveness (collapsed across label). Here, the grey shading signifies significant temperature differences between attractive and unattractive faces. A two-sided paired cluster-based permutation test identified a significant cluster, highlighted in dark grey. This cluster indicates a sustained difference in facial temperature between conditions, corrected for multiple comparisons across all time points. In all panels, where applicable, error bars represent the standard error of the mean.

### Mean subjective attractiveness rating

Figures 2a and 2b illustrate the mean attractiveness rating as a function of the autism label and facial attractiveness for male and female participants, respectively. We conducted a repeated measures ANOVA on the mean attractiveness rating with facial attractiveness (attractive versus unattractive faces) and autism label as within-subject factors and sex as between-subject factor. Shapiro–Wilk tests indicated that all conditions were normally distributed (all *p*-values ≥.954), confirming that the assumptions for the ANOVA were met. Even though the labels were randomly determined on every trial, the ANOVA revealed a significant effect of the autism label, *F*(1, 38) = 6.87, *p* =.013, partial *η*^2^ =.153, such that faces preceded by an autism label (M = 36.1) were rated as less attractive than those preceded by no such label (M = 37.2).However, the sex × label interaction was significant, *F*(1, 38) = 7.73, *p* =.008, partial *η*^2^ =.169, indicating that the effect of the autism label depended on participants’ sex. Male participants rated the faces as less attractive when they were preceded by an autism label (M = 37.7) than when preceded by a no-autism label (M = 40.1), *t*(11) = 3.02, *p* =.012 (two-tailed *t*-tests). In contrast, the label effect was not significant for female participants, as attractiveness ratings were nearly identical for faces preceded by an autism label (M = 34.4) and those preceded by a no-autism label (M = 34.3), *t*(27) = 0.15, *p* =.882. The ANOVA yielded a significant facial attractiveness × autism label interaction, *F*(1, 38) = 6.40, *p* =.016, partial *η*^2^ =.144. Follow-up two-tailed *t*-tests showed that the label effect was significant for attractive faces, *t*(39) = 2.40, *p* =.021, but not for unattractive faces, *t*(39) = 0.94, *p* =.355, possibly reflecting a floor effect in the latter condition. All other effects failed to reach significance, all *F*(1, 38) values ≤ 2.66, all *p* values ≥.111.

To assess the generalizability of our attractiveness ratings, we correlated the mean attractiveness ratings of each face across participants obtained in the present study with the normative ratings provided by the original face database (Ma, Correll, & Wittenbrink, 2015). The two measures were strongly and positively correlated, Pearson *r* =.65, *p* <.0001, indicating that participants’ judgments closely aligned with large-scale normative evaluations of facial attractiveness. This confirms that the observed label effects are not driven by idiosyncratic rating behavior but reflect shared standards of facial attractiveness.

To ensure the observed label effect was not confounded by face assignment, we tested whether any of the 48 faces were labeled autistic more or less often than expected by chance (*p* = 0.5) using binomial tests, separately for male and female images, with FDR correction. No face deviated from the expected distribution (all FDR corrected *p*-values ≥.509), confirming that randomization successfully balanced faces across label conditions.

### Mean nose temperature change

Figures 2e and 2f show the mean change in nose temperature as a function of the autism label and facial attractiveness for male and female participants, respectively. A repeated-measures ANOVA was conducted on the mean nose temperature change, with autism label and facial attractiveness as within-subject factors and sex as a between-subjects factor. Shapiro–Wilk tests indicated that all conditions were normally distributed (all *p*-values ≥.877). The ANOVA revealed a significant main effect of facial attractiveness, *F*(1, 38) = 5.34, *p* =.026, partial *η*^2^ =.123, such that attractive faces elicited a greater increase in nose temperature (M = 0.014°C) than unattractive faces (M = −0.019 °C). There was also a significant main effect of sex, *F*(1, 38)= 5.48, *p* =.025, partial *η*^2^ =.126, with female participants showing a small increase in nasal temperature (M = 0.008 °C) and male participants showing a decrease (M = −0.026 °C). No other effects were significant, all *F* < 1, indicating that the autism label did not influence mean nose temperature change.

Subsequently, we examined the temporal dynamics of the facial attractiveness effect. Figure 2g shows the mean nose temperature change as a function of time since image onset (collapsed across label conditions). To assess differences between attractive and unattractive faces across the full time series (120 samples), we conducted a two-sided paired cluster-based permutation test on sample-wise *t*-tests (Maris & Oostenveld, 2007). Clusters were formed using an uncorrected threshold of *p* <.05, and significance was evaluated against a permutation-derived null distribution of the maximum cluster statistic using 5,000 permutations. This approach controls for multiple comparisons while accounting for the temporal dependency of the data. Cluster statistics were computed as the within each contiguous cluster of samples exceeding the threshold. Sample-wise comparisons indicated emerging differences between conditions from approximately 4 s post-stimulus (light grey shading in Figure 2g). Critically, the permutation analysis identified a significant cluster of elevated nasal temperature for attractive relative to unattractive faces between 10.0 and 14.7 s after stimulus onset (samples 75–110; cluster-level *p* =.012; dark grey shading in Figure 2g), reflecting a sustained attractiveness-related modulation of facial temperature.

Next, we examined whether changes in mean nose temperature predicted the perceived attractiveness of the female and male images across participants. Shapiro–Wilk tests indicated that the variables were normally distributed for both female images (*p* =.070) and male images (*p* =.095), confirming that Pearson correlations were appropriate. As shown in Figure 2c, there was a significant positive correlation between nose temperature change and attractiveness ratings for female faces (Pearson’s *r* =.297, *p* =.040), indicating that greater temperature increases were associated with higher attractiveness ratings. For male faces, the correlation was also positive but did not reach significance (*r* =.132, *p* =.370; see Figure 2d).

### Empathy

The mean BES total score was 82.2 (range: 64–97). We examined whether the mean nose temperature change and the subjective attractiveness rating correlated with the different BES scores. Shapiro–Wilk tests showed that the BES total and subscales were normally distributed (all *p*-values ≥.057) and subjective attractiveness ratings were normally distributed (*p* =.356), whereas the mean nose temperature change was not (*p* =.013). Therefore, Spearman correlations were used for analyses involving nose temperature. Nose temperature change correlated positively with the BES total score (Spearman’s *ρ* =.568, *p* <.001), the BES cognitive subscale (*ρ* =.469, *p* =.002), and the BES affective subscale (*ρ* =.502, *p* <.001), indicating that greater nose temperature increases were associated with higher empathy scores (see Figures 3a–c). Pearson correlations showed the same pattern of results (all *p*-values ≤.016), confirming the robustness of these findings. As shown in Figures 3d–f, subjective attractiveness ratings did not correlate with the BES total score (Pearson’s *r* =–.219, *p* =.176), the cognitive subscale (*r* = –.270, *p* =.092), or the affective subscale (*r* = –.135, *p* =.405). Together, these findings suggest that individual differences in empathic ability are reflected in physiological responses to social stimuli, as indicated by changes in nose temperature, but not in the explicit attractiveness ratings of faces.

**Figure 3.**
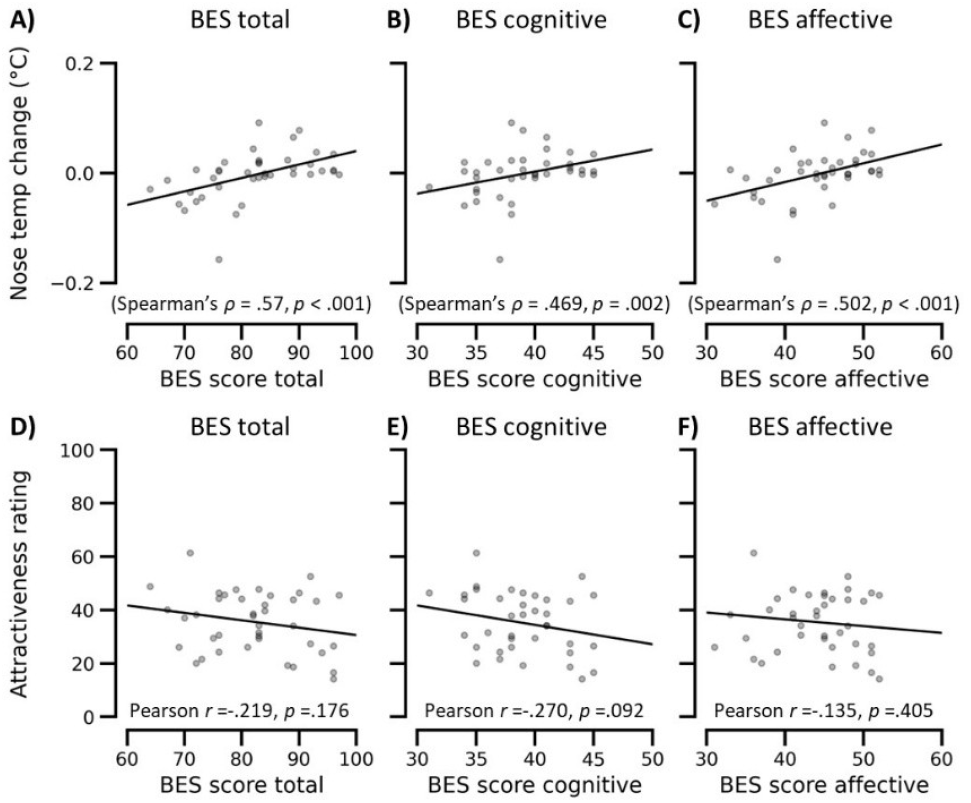
A-C) Mean nose temperature change as a function of the basic empathy scale (BES) scores (from left to right: total BES, cognitive and affective score). D-F) Mean attractiveness rating as a function of the BES scores.

### Socials distance to, and familiarity with, autism

The mean social distance score (i.e., stigma) was 19.4 (range: 13–24), and the mean familiarity score (i.e., level of contact with autistic individuals) was 7.15 (range: 2–11). We next examined whether these measures predicted either the subjective effect of the autism label (attractiveness rating for faces with an autism label minus attractiveness rating for faces without a label) or the objective effect (nose temperature change for faces with an autism label minus nose temperature change for faces without a label). Because the questionnaire scores were not normally distributed (Shapiro–Wilk tests, all *p* ≤.035), non-parametric correlations were used. Social distance was not significantly associated with the subjective rating difference (Spearman’s *ρ* = –.206, *p* =.203) nor with the objective temperature difference (*ρ* =.186, *p*=.250). Likewise, familiarity with autism did not correlate with the subjective rating difference (*ρ* =.061, *p* =.707) or the objective temperature difference (*ρ* =.010, *p* =.952). Thus, neither stigma nor familiarity with autism appeared to influence participants’ behavioural or physiological responses to the autism label.

## Discussion

The present study investigated whether autism-related stigma biases facial attractiveness evaluations and whether such biases are reflected solely in explicit judgments or also in automatic physiological responses measured using fITI. The results reveal a clear dissociation between subjective and physiological measures. While autism labels systematically influenced explicit attractiveness ratings, particularly among male participants and for highly attractive faces, facial thermal responses were unaffected by diagnostic labels. Crucially, we show that perceived facial attractiveness reliably predicts thermal responses, with nose temperature tracking attractiveness such that attractive faces elicited greater and sustained temperature increases relative to unattractive faces. These findings suggest that autism-related labels shape consciously reported evaluations, whereas autonomic physiological responses index more fundamental perceptual and affective processing of facial attractiveness, largely independent of socially constructed diagnostic information.

Our findings show that diagnostic labels altered how faces were judged explicitly, but not how they were processed physiologically. This pattern contrasts with the results of Brosnan and Gavin (2021) and Gavin and colleagues (2019), who reported that an autism label increased perceived attractiveness when female participants evaluated the attractiveness of a single male individual on a dating profile page. In the present study, however, we observed the opposite pattern: male participants rated faces as less attractive when they were preceded by an autism label, whereas no such effect was observed for female participants. This divergence suggests that the impact of autism labels on attractiveness judgments is highly context-dependent and shaped by characteristics of both the observer and the evaluative task, including participant sex, stimulus format, and repeated exposure to multiple faces. Notably, the label effect in our study was restricted to faces that were rated as attractive, whereas unattractive faces showed little modulation by diagnostic labels, likely reflecting limited room for further devaluation given their low attractiveness ratings. Together, these findings indicate that autism labels function as social meaning cues that selectively reshape explicit evaluative judgments rather than perceptual encoding of facial information.

The present study provides the first evidence that facial attractiveness systematically modulates facial thermal responses. Specifically, nose temperature increased more for attractive than for unattractive faces, with this effect emerging approximately 10–15 seconds after stimulus onset. Importantly, this does not imply that nose temperature functions as a “beauty detector.” Rather, we propose that facial attractiveness engages the autonomic nervous system by virtue of its affective and motivational significance. Changes in nasal temperature are known to reflect vasodilation and vasoconstriction driven by sympathetic and parasympathetic control of cutaneous blood flow (Ioannou et al., 2014; Pavlidis et al., 2002), processes unfolding over several seconds, consistent with the delayed temporal profile observed here. Converging evidence shows that nose temperature covaries with other indices of autonomic arousal, such as skin conductance (Kosonogov et al., 2017; Pavlidis et al., 2012), and responds to a broad range of affective stimuli, including stress (Engert et al., 2014), joy (Nakanishi & Imai-Matsumura, 2008), sexual arousal (Merla & Romani, 2007) and more (Ioannou et al., 2014). From this perspective, the observed thermal modulation reflects domain-general affective engagement rather than stimulus-specific aesthetic processing. Consistent with this interpretation, prior work has shown that nasal temperature increases when individuals think about their romantic partner compared to thinking about someone else’s relationship or when watching a video message of one’s own loved one compared to watching a video of another person’s loved one (Sprengel et al., 2025), indicating that nose temperature is sensitive to affiliative and attachment-related states rather than perceptual features alone (Cannas Aghedu et al., 2022). As a fully non-invasive and contact-free technique, fITI may provide a powerful tool for studying implicit affective responses to socially relevant stimuli beyond faces, including music, food, or other pleasant and unpleasant experiences, offering a complementary window into evaluative processes largely inaccessible to self-report.

Individual differences in empathy robustly predicted physiological responses to faces, while remaining unrelated to explicit attractiveness judgments. Participants with higher empathy scores showed larger changes in nose temperature when viewing faces, yet their attractiveness ratings did not differ systematically from those of less empathic individuals. These findings suggest that facial thermal responses are sensitive to the observer’s affective disposition, whereas explicit ratings primarily reflect consciously reported, and potentially socially filtered, evaluations. In other words, nasal temperature appears to index how strongly an observer *feels* in response to a face, whereas attractiveness ratings reflect what observers *choose to report*. This interpretation aligns with a broader literature showing that empathic traits selectively modulate autonomic and neurophysiological responses to social stimuli. For example, higher empathy has been associated with stronger skin conductance responses when observing others in pain (Hein et al., 2011), and greater pupil dilation to emotional facial expressions (Geangu et al., 2011). Together with the present findings, this body of work supports the view that empathy amplifies physiological sensitivity to socially and emotionally salient stimuli without necessarily altering explicit evaluative judgments. Taken together, the observed empathy–temperature relationship reinforces the interpretation of nasal thermal responses as a marker of affective engagement rather than deliberate aesthetic appraisal, further underscoring the functional dissociation between subjective reports and autonomic indices of social perception.

The present study also has several methodological limitations. To ensure stable thermal measurements, participants’ head position was constrained using a chinrest and trials with excessive movement were excluded using a simple algorithm. Future work could leverage automated facial landmark detection methods that track regions of interest, such as the tip of the nose, even under moderate head motion (e.g., Sonkusare et al., 2019), enabling fITI measurements under more naturalistic and unconstrained conditions. Importantly, despite these constraints, we observed reliable and systematic modulations of nasal temperature as a function of facial attractiveness, underscoring the robustness of the present effects. A second limitation concerns the use of highly controlled, static facial stimuli. While this approach affords strong experimental control, it may elicit weaker emotional and motivational engagement than faces encountered in real-world social interactions. Moreover, the stimulus set was skewed toward relatively unattractive faces, limiting variability at the upper end of the attractiveness distribution. Both factors likely render the observed effects conservative. Future studies using more diverse and ecologically valid stimuli, such as dynamic, expressive faces spanning a broader attractiveness range, may yield stronger and more differentiated physiological responses, further clarifying how affective engagement shapes social evaluation.

In sum, the present study reveals a clear dissociation between subjective and physiological responses to facial stimuli: autism labels influenced explicit attractiveness judgments but did not alter autonomic engagement, as indexed by nasal temperature. This is particularly relevant given evidence that autistic individuals, compared to non-autistic peers, often prefer online dating as a social and romantic medium (M. Neumann et al., 2024; Roth & Gillis, 2015). While Brosnan and Gavin (2021) suggested that explicitly disclosing an autism diagnosis on a dating profile could enhance perceived attractiveness, our results indicate that such effects are modest and context-dependent: male participants rated faces with an autism label slightly lower (∼2.4%) than neutrally labeled faces, whereas female participants were unaffected. Crucially, the absence of physiological modulation suggests that these label effects do not reflect deeper affective responses, implying that disclosure is unlikely to provoke negative emotional reactions. Taken together, our findings demonstrate that while social labels shape explicit judgments, the underlying autonomic system responds to facial features themselves, so, in matters of facial attractiveness, the nose knows.

## Acknowledgement

We would like to thank Julie Al, Florentina Boekelaar, Madelief Flentge, and Sanne van der Gugten for recruiting participants and for collecting the data.

